# Transposon fusion gave birth to *Fem*, *Bombyx mori* female determining gene

**DOI:** 10.1101/2024.01.09.574859

**Authors:** Jung Lee, Toshiaki Fujimoto, Katsushi Yamaguchi, Shuji Shigenobu, Ken Sahara, Atsushi Toyoda, Toru Shimada

## Abstract

*Fem* is a W-linked piRNA precursor, and *Fem* piRNA is a master gene of female determination in *Bombyx mori*. Since *Fem* has significantly low similarity to any known sequences and the origin of *Fem* remains unclear, two hypotheses have been proposed for the origin of *Fem*. The first one is that *Fem* is an allele of *Masc,* while the second one is that *Fem* arose by transposition of *Masc*. To draw a solid conclusion, we determined the W chromosome sequences of *B. mori* and a closely relative bombycid species of *Trilocha varians* with a *Fem*-independent sex determination system. Comprehensive genome comparison led us to the “third” hypothesis: *Fem* is a chimeric sequence of transposons. Although we still cannot completely exclude the first hypothesis, at least a large portion of the parts other than the 44-bp *Masc* similarity region is derived from transposons, and even the 44-bp region could correspond to the boundary of the two transposons, gypsy and satellite DNA.

## Introduction

*Fem* is a W-linked female sex-determining gene in *Bombyx mori*. *Fem* encodes a piRNA precursor of 767 bases [1]. The sequence of the piRNA-producing region in *Fem* is partially complementary to the coding sequence of *Masc*, a Z-linked male sex-determining gene. *Fem* piRNAs generated from *Fem* mRNAs recognize *Masc* mRNAs based on sequence homology, and Piwi protein forming a complex with *Fem* piRNA disrupts *Masc* mRNAs, resulting in female differentiation in W chromosome carriers. The issue we attempt to address in this paper is the origin of *Fem* because sequences outside the piRNA-producing region in *Fem* have no homology to any known genes. So far, two hypotheses have been proposed for the origin of *Fem*, as described below. One is that *Fem* was originally an allele of *Masc*. W and Z chromosomes were originally homologous chromosomes and the W-linked allele diverged to the piRNA- producing gene as sex chromosome heteromorphism proceeded. The other scenario is that *Masc* was transposed to the W chromosome, which gave rise to *Fem*. In the case of the diamondback moth (*Plutella xylostella*), the researchers insisted that the sequences of *P. xylostella Fem* (*Pxyfem*) retain the traces of the retroposition of *Masc* [2]. Both of the two hypotheses are not essentially different in that they assume *Fem* originated from *Masc*. However, we could not conclude which hypothesis applies to the case of *B. mori* because the W chromosome of *B. mori* is not determined.

Recent studies suggested that the W chromosomes of butterflies have multiple origins. The collinearity between the W and Z chromosomes of two *Pieris* butterflies, namely *Pieris manni* and *Pieris rapae*, exists, suggesting that the W chromosomes of *P. mannii* and *P. rapae* have evolved from Z chromosomes [3] while the W chromosome of another butterfly species, *Dryas iulia*, is derived from the B chromosome [4]. Lepidopteran W chromosomes tend to accumulate retrotransposons and often have complex structures [4,5], and *B. mori* represents a prime example: because the nested structures of the W- linked retrotransposons were of considerable degree [6,7], chromosome-scale assembly of the W chromosome of *B. mori* has been considered impossible. For this reason, in the genome projects organized in the past [8–10], male individuals have been used as the donor of genomic DNA. We attempted to determine and scaffold the W chromosome sequences of *B. mori* with long-read sequencing and optical genome mapping technology to elucidate the origins of *Fem* and W chromosomes. We therefore sequenced the W chromosome of *B. mori* using PacBio Hifi reads and Bionano Saphyr, two of the available technologies which have advantages in assembling hyper-repetitive sequences. As a comparison, in addition to the W chromosome of *B. mori*, we sequenced the W chromosome of the *T. varians* closely related species to *B. mori*, which was expected not to possess *Fem*. As a result, we succeeded in constructing chromosome-scale genome assemblies. Comparative W chromosome analysis led us to suggest the third hypothesis that *B. mori Fem* was not an allele of *Masc*. A fusion event of multiple transposons and the sequence that arose at the boundary of the fusion happened to have a complementary sequence to *Masc*, which took on a function and became a *Fem*.

## Results& Discussion

### Genome assembly and completeness assessment

We constructed chromosome-scale assemblies of the female genomes of *B. mori*. For comparison, the female genome of *T. varians*, which was expected not to possess *Fem* [11], was also determined (Fig. 1A). We successfully scaffolded all chromosomes into a single sequence for both species (Table 1; S1 Table). Of the 5286 lepidopteran BUSCOs, 98.6% and 98.7% were found to be complete within our *B. mori* assembly (BmF-NBRP_2.0) and *T. varians* assembly (TvF-NBRP_1.0), respectively (S2 Table). Mapping rates of long-read to the assemblies were 99.99% (BmF-NBRP_2.0) and 99.87% (TvF- NBRP_1.0), respectively. Moreover, regarding the assembly of *B. mori*, autosomal and Z-linked regions of both assemblies were consistent with the latest male assembly (S1 Fig) [12]. Respective self-alignments of both assemblies yielded numerous small alignments on W chromosomes, suggesting that the W-linked sequences were highly repetitive, as expected (S1 Fig). Since there is no genome assembly of *T. varians*, an extra assembly assessment was carried out by BAC-FISH mapping. Each BAC-FISH signal reflected a positional relationship on the assembly (S2 Fig). Based on these results, we concluded that the quality of both assemblies is comparable to that of previous assemblies, at least for the Z chromosome and autosomes.

**Figure 1.**
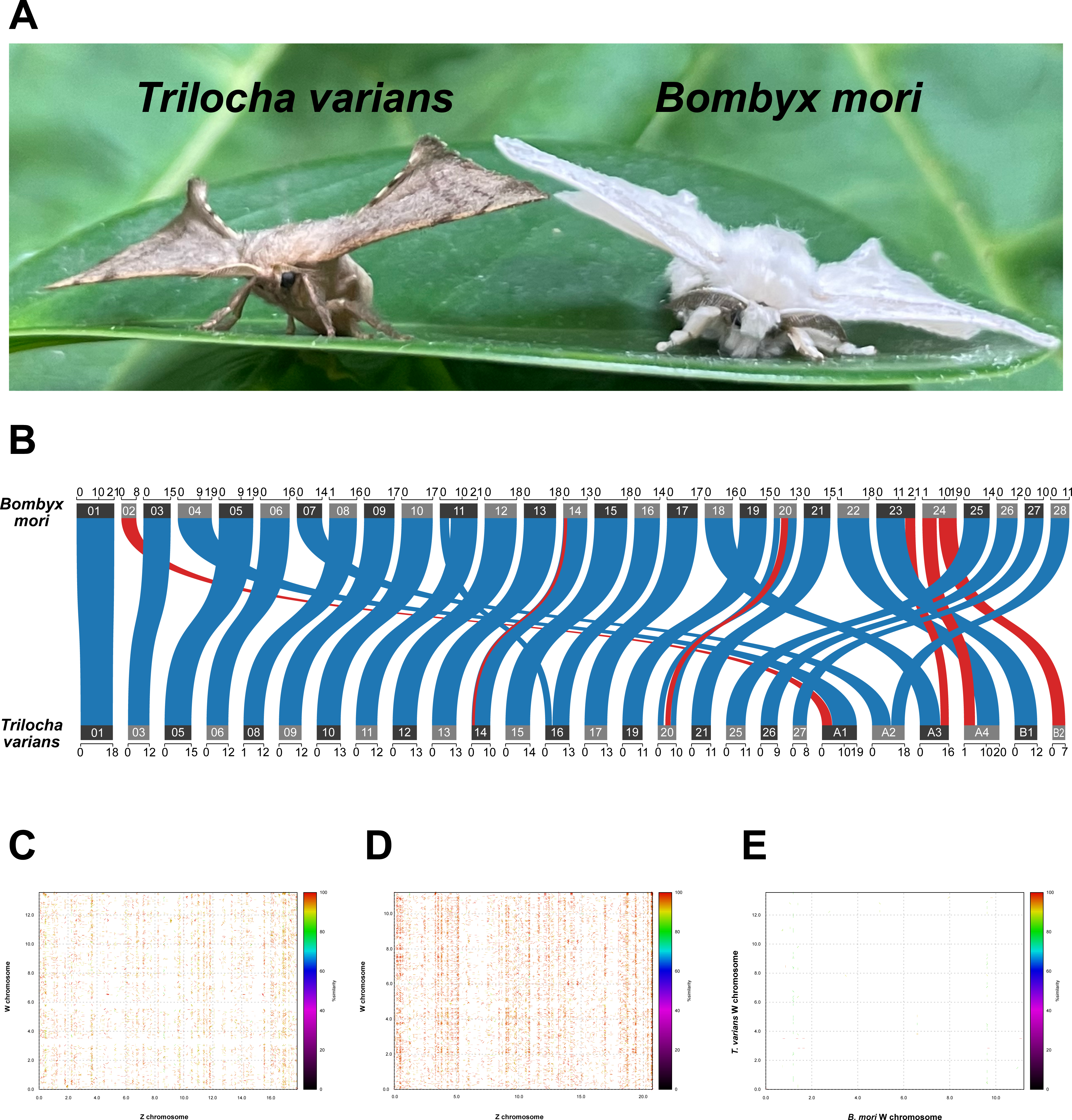
Female genome assemblies of *T. varians* and *B. mori*. **A.** Adult female moths of *T. varians* (left) and *B. mori* (right). **B.** The genomic loci of common single-copy orthologs (SCO) between *B. mori* and *T. varians* were linked. Blue links indicate two SCOs have the same direction, while red links indicate two SCOs have inverted directions. **C.** Alignment of the Z chromosome (X-axis) and W (Y-axis) in *T. varians*. **D.** Alignment of the Z chromosome (X-axis) and W (Y-axis) in *B. mori*. **E.** Alignment of the W chromosome of *B. mori* (X-axis) and W chromosome of *T.varians* (Y-axis). **B-E.** The numbers alongside each axis represent the genomic coordinates in the megabase scale.

**Table 1.**
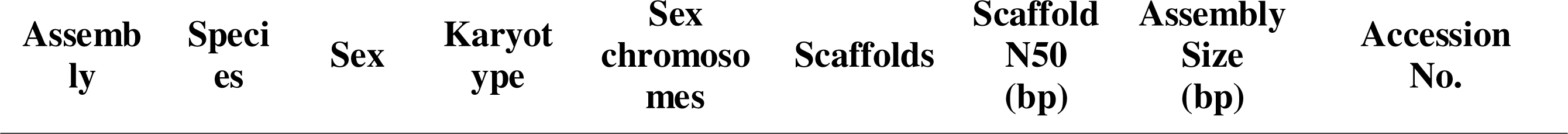
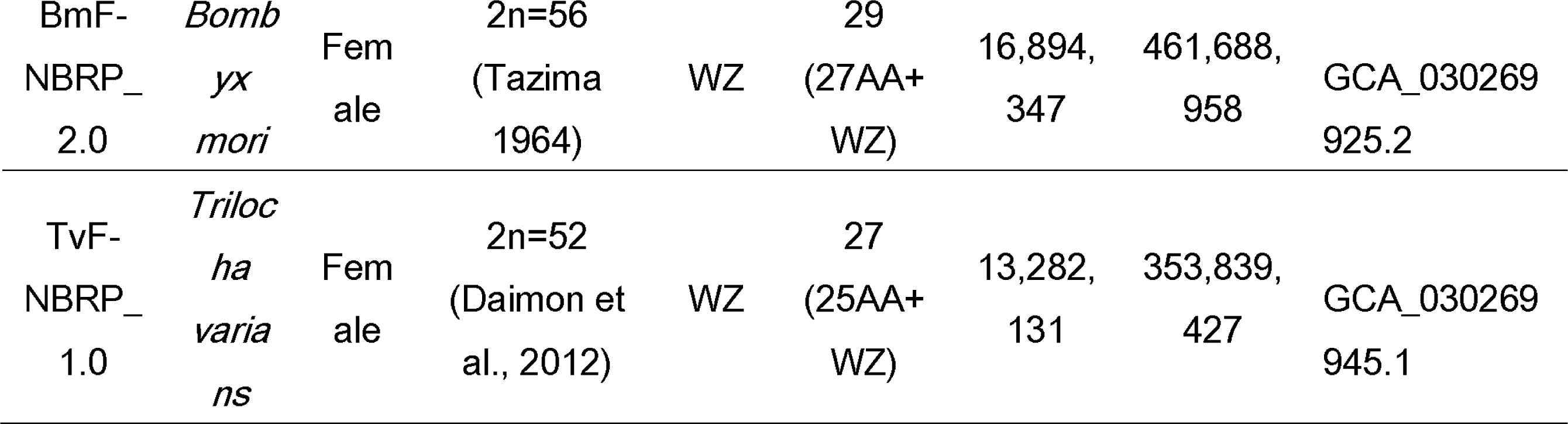
Overview of statistics of two assemblies.

| Assembly | Species | Sex | Karyotype | Sex chromosomes | Scaffolds | Scaffold N50 (bp) | Assembly Size (bp) | Accession No. |
| --- | --- | --- | --- | --- | --- | --- | --- | --- |
| BmF-<br>NBRP_<br>2.0 | *Bomb*<br>*yx*<br>*mori* | Fem<br>ale | 2n=56<br>(Tazima<br>1964) | WZ | 29<br>(27AA+<br>WZ) | 16,894,<br>347 | 461,688,<br>958 | GCA_030269<br>925.2 |
| TvF-<br>NBRP_<br>1.0 | *Triloc*<br>*ha*<br>*varia*<br>*ns* | Fem<br>ale | 2n=52<br>(Daimon et<br>al., 2012) | WZ | 27<br>(25AA+<br>WZ) | 13,282,<br>131 | 353,839,<br>427 | GCA_030269<br>945.1 |

For the *B. mori* W chromosome, we also carried out BAC-FISH mapping, but all of the selected BAC clones painted the entire chromosome or the end of a non-W chromosome (S2 Fig). It was nearly impossible to select region-specific BAC clones because approximately 90% of the W chromosome of *B. mori* is occupied by repetitive sequences, as previously predicted (S3 Fig). Because the structure of the W chromosome in *T. varians* is relatively simpler than that of *B. mori* (S2 and S3 Figs), it was expected that selection of position-specific BAC clones was not impossible, but the more fundamental problem arose: the impossibility of chromosome specimen preparation using female specimens. Single copy orthologs (SCO) anchored analysis revealed overall synteny of autosomes and Z chromosomes was well conserved (Fig. 1B) even though chromosome fusion and fragmentation had occurred several times after the divergence of the common ancestor of the two species. However, the assembled W chromosome sequences of *B. mori* and *T. varians* did not share any SCOs, and no synteny or homology could be detected at the scale of nucleotide sequence (Fig. 1C–E; S1 Fig). Therefore, we concluded that it was impossible to perform assembly assessment of the W chromosome by experimental methods.

### W chromosome content and synteny analysis

RepeatMasker annotated more repetitive elements on the W chromosomes than autosomes or Z chromosomes in both species (S3 Fig). As previously predicted [6,7], an extensively large fraction of the *B*. *mori* W chromosome, namely 87.74% of the total length, was occupied by repetitive sequences, which could explain the inability to detect sequence homology between Z and W chromosomes. Then, we performed embryonic RNA-seq and Iso-seq to identify orthologs shared by Z and W chromosomes. While five Z-linked genes and one W-linked gene, whose homologs were linked to the counterpart chromosomes, were identified in the embryonic transcriptome data of *T. varians* (S4 Fig). Although the five pairs of Z-W homologs found in *T. varians* could not be located on the *B. mori* W chromosome, only another pair of Z-W homologs was found in the *B. mori* (S3 Table), probably due to the recent explosive proliferation of transposons (S5 Fig), which could have destroyed the synteny on the W chromosome. The collinearity between the Z and W chromosomes of *T. varians* was subtle but undoubtedly present, which not only illustrates the W chromosome of *T. varians* was derived from Z chromosome but paradoxically provides assurance about the accuracy of the W chromosome assembly of *T. varians*. *T. varians* have also recently experienced a transposon explosion, but it was to a lesser extent (S5 Fig), and hence the collinearity between Z and W chromosomes was detected. Although only one pair of Z-W homologs was not enough to define synteny between sex chromosomes in *B. mori*, this result supports the Z chromosome origin of the W chromosome in *B. mori*.

In some species, W chromosomes were presumed to have derived from the B chromosome [4,13]. The primary basis of this kind of non-canonical “B chromosome” theory of W chromosome evolution is that gene repertoires of Z chromosomes are extensively conserved among lepidopteran insects. This fact indirectly refutes the canonical theories, such as the sex chromosome turnover or neo-sex chromosome theory. However, as shown in S4 Fig, examining individual genes would find traces of the Z chromosome in the W chromosome. The accumulation of chromosome-scale genome assemblies and transcriptome information would improve our understanding of the origin of the W chromosome.

### *Fem* is absent in the W chromosome of *T. varians*

Although W and Z chromosomes are homologous to each other in *T. varians*, we could hardly find the *Masc* homologue on the *T. varians* W chromosome (Fig. 4). To confirm the absence of W-linked *Masc* homologue or *Masc*-targeted piRNA-producing gene (putative *Fem* orthologue), we performed deep sequencing of embryonic piRNA-libraries of *T. varians* and searched piRNA reads complementary to *Masc*. As expected, we did not identify piRNA reads with complementary sequences to *TvMasc,* while *B. mori* and *B. mandarina* have *Fem* transcripts and piRNA reads with complementary sequences to *Masc* (S6 Fig). Given the previous report [11], we concluded that *T. varians* lacks W-linked *Masc* homolog (putative *Fem* ortholog) and adopts a piRNA-independent sex-determination system. As in *Samia cynthia ricini*, a lepidopteran species with a Z0/ZZ sex chromosome system, Z: A ratio might be essential in the sex determination of *T. varians* [14].

### *Fem* of both *B. mori* and *B. mandarina* are not derived from the retrotransposition of *Masc*

The *Fem* of *P. xylostella* (*Pxyfem*) have arisen from retrotransposition of *Masc* to the W chromosome [2]. piRNA producing region of *Pxyfem* is homologous to exon-exon boundaries of exon 5 and 6 of *PxyMasc* whereas that of *B. mori Fem* corresponds to exon 9. Therefore, even if *BmoFem* was also derived from retrotransposition of *Masc*, *BmoFem* and *Pxyfem* have different origins. Our assembly of the *B*. *mori* W chromosome has 129 copies of *Fem* (Fig. 2A). *Fem* copies were not dispersed randomly on the W chromosome. Rather, they were concentrated in multiple copies, forming clusters. We located 11 *Fem* clusters on the W chromosome in total (Fig. 2A). FISH mapping also confirmed some clusters of *Fem* (S7 Fig), suggesting that the *Fem* clusters were not the side product of misassembly. *Fem* copies within a cluster have the same direction with no exception, which suggested multiplication of *Fem* was not achieved through retrotransposition (Fig. 2B). Although there are traces of retrocopy at the 3’ ends of *Pxyfem*, we could not find any traces of retrotransposition in *BmoFem* transcripts or in the adjacent genomic regions of *BmoFem*s (Fig. 2A). Samuel et al. (2022) described the W chromosome as “transposable element graveyard” as if the entire W chromosome is a huge piRNA cluster (piC). They may assume a scenario in which the retrotransposition of *Masc* to such “transposable element graveyards” triggered the production of *Pxyfem* piRNAs with complementary sequences to *Masc*. The piRNA cluster “*torimochi*” in BmN cells, a cultured cell line of *B. mori*, is known to produce piRNAs with complementary sequence of an inserted exogenous gene [15]. However, in our assembly, none of the above-mentioned 11 *Fem* clusters overlaps with the position of the W-linked piRNA cluster (Fig. 2A). Taken together, the “retrotransposition origin” seems to be a less plausible assumption in *B. mori*.

**Figure 2.**
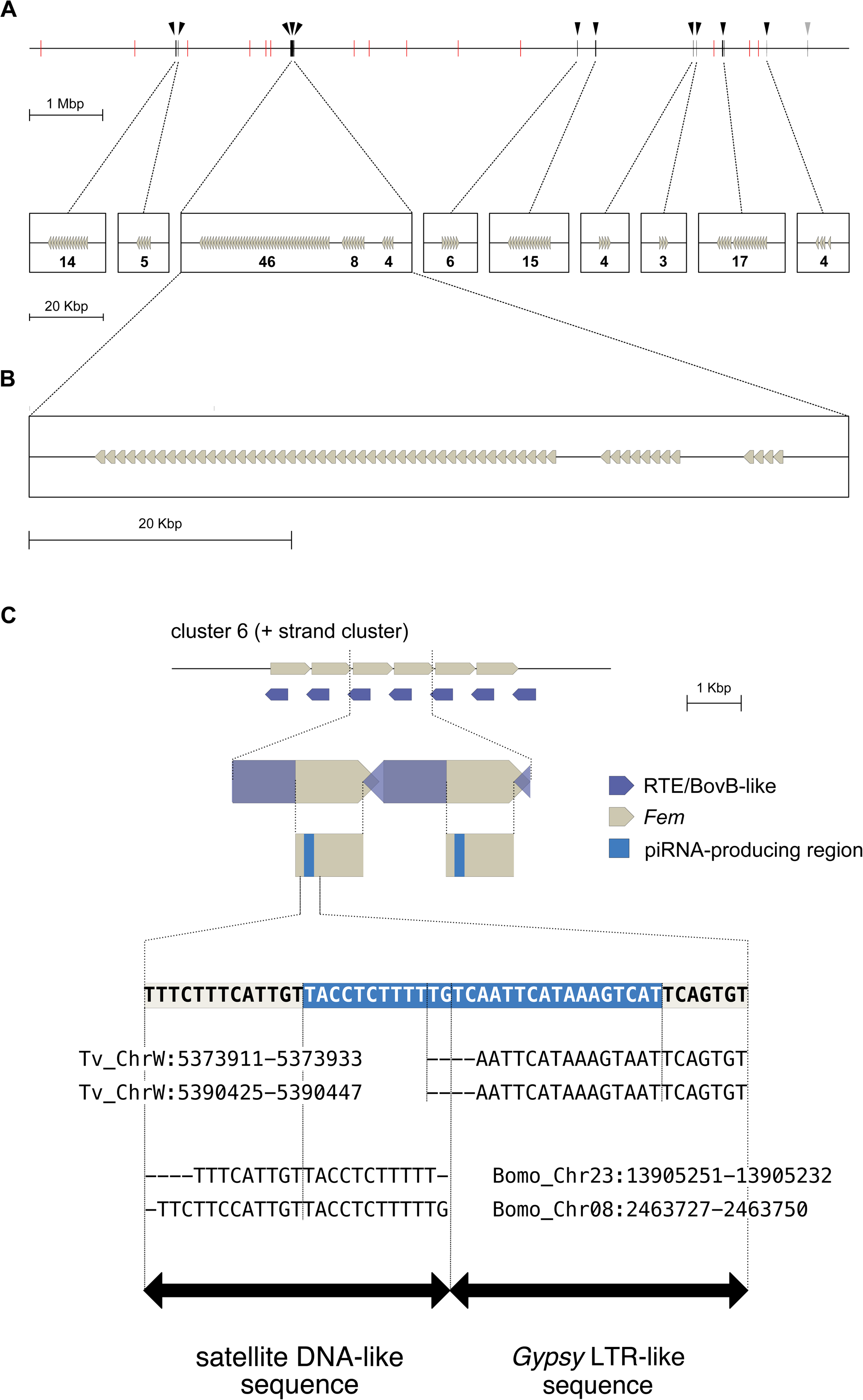
The schematic diagrams of *Fem* clusters and *Fem* structure. **A.** *Fem* distribution map on BmW. Arrowheads indicate *Fem* clusters. We defined a “*Fem* cluster” as follows: a *Fem* cluster should have more than three copies of *Fem* within one Kbp or shorter proximity. According to this definition, there were 11 *Fem* clusters on BmW. The rightmost arrowhead indicates the presence of 4 copies of *Fem*, but the distance between the copies exceeds 1 kbp, so they are not defined as a “cluster”. The upper scale bar indicates 1 megabase, and the lower one does 20 kilobases. Red vertical bars are the position of W-linked piRNA clusters. **B.** Orientations of *Fem* copies within the same *Fem* clusters. All copies have the same orientation in a cluster with no exceptions. The most massive cluster, cluster 3, and its neighbours, clusters 4 and 5, are shown as examples. All copies illustrated here are coded on the positive strand. The scale bar indicates 20 kilobases. **C.** The elements comprise a *Fem* sequence. Both terminals of a *Fem* show a high similarity to the RTE-type LINE BovB. The first half of the inner *Fem* piRNA-producing has a satDNA-like sequence, while the second half shows a similarity to the LTR of Gypsy. The results of the multiple alignments of the transposons, estimated to be the ancestors of *Fem,* were visualised in the sequence logo and placed under the corresponding *Fem* sequence. The scale bar indicates 1 kilobase.

### *Fem* is a chimeric gene of transposons

The simplest hypothesis on the origin of *Fem* is that *Fem* was a diverged allele of *Masc*. Since 44 bp-length around the *Fem* piRNA-producing region shows considerable similarity to exon 9 of *Masc* (we call this region the “high-similarity region (HSR)” afterward), it is impossible to completely deny this hypothesis. However, some questions remain about this hypothesis. Besides *Fem*, *angel* is the only gene whose homologs are located on both W and Z chromosomes. The Z-linked homolog of *angel* (*Z_angel*) has one copy, but there are two identical copies on the W chromosome (*W_angel1* and *W_angel2*). Although transcripts of neither *W_angel1* nor *W_angel2* have been identified probably due to pseudogenization, the ORFs of *W_angel1* and *W_angel2* are almost identical to CDS of *Z_angel* except for some mutations. When we mapped small RNA reads derived from early embryos to the putative CDS of *W_angel1* and *W_angel2*, some reads were mapped to the CDS, but did not show 1U-bias (S8 Fig). This observation points out that the gene on an excessively diverged heterosome does not necessarily start producing piRNA. If *Fem* is the allele of *Masc*, how did *Fem* acquire the ability to produce piRNA?

Whether the HSR is derived from *Masc* or not, approximately half of *Fem* is unambiguously derived from the transposon. Regarding the outside regions of the piRNA-producing region, it shows significant similarity to BovB, an RTE-type LINE (Fig. 2B and S9 Fig). BovB elements span the *Fem*- *Fem* boundaries in the reverse direction (Fig. 2C). In other words, the 5’ and 3’ ends of *Fem* correspond to the 3’ and 5’ ends of BovB, respectively. A typical BovB sequence lacks ORF1 and has only ORF2. The BovB sequences between *Fem* copies also have disrupted ORF2 (S9 Fig). Since all 129 *Fem* copies are associated with the BovB-derived sequence, *Fem* increased its copy number after the fusion of BovB to the ancestral *Fem* sequence (proto-*Fem*). In addition, we realized that *Fem* produces different piRNAs other than *Fem* piRNA (S10 Fig). Interestingly, such piRNAs have 10-base pair overlap with small RNA reads derived from TE transcripts (asmbl_70383 and asmbl_39053), suggesting these piRNAs constitute the ping-pong cycle. Thus, it is more plausible to consider that this region originated from transposon rather than *Masc*.

Then, was the HSR of *Fem* really derived from *Masc*? Performing BLAST search using 50 bp, including the HSR of *Fem* as a query, we yielded the alignments to *Fem* and *Masc* with the lowest e- values, but also to autosomal sequences (S4 Table). Such alignments are yielded not only in the genome assemblies of *B. mori* and *B. mandarina* but also in phylogenetically distant species, *Antheraea yamamai* (S4 Table). These BLAST results led us to suspect that even HSR could be derived from transposons. First, we focused on the 5’ half of the HSR. BLASTN searches with 5’ half of the *Fem* piRNA-producing region as query on the genomes of *B. mandarina* and *B. mori* yielded the alignments with e-value<0.05 to chromosome 8 of both species. A close examination located this site at a terminal region of the tandem repeat (TR) cluster (S11 Fig). This region contains two clusters of TRs (TRC1 and TRC2; S11 Fig), and each cluster comprises highly similar repeat units (motif1 and motif2; S11 Fig). In the homologous region on chromosome 8 of *B. mandarina,* there are three TRCs, while *T. varians* has no TRs in the same region (S12 Fig). Because the 5’ sequence of the *Fem* piRNA-producing region showed the highest similarity to the edges of the TRC array in both *B. mandarina* and *B. mori* (S11 and S12 Figs), we estimated that each repeat unit at the 3’ edge of the cluster of *B. mori* and *B. mandarina* are homologous. The size of a single repeat unit (approximately 150 bp) rationalizes these TRs into satellite DNA (satDNA). Many studies have provided examples of transposable sequences being one of the primary sources for the birth of TR, e.g., microsatellites, minisatellites, and satellite DNA [16–18]. Since this TR cluster functions as a piRNA cluster (S11 Fig) and significantly similar sequences to the 5’ sequence of the *Fem* piRNA-producing region were identified in multiple chromosomes (i.e., chromosome 23 in addition to chromosome 8; S4 Table), we postulated that satDNA (and the 5’ sequence of the *Fem* piRNA-producing region) was derived from the transposable element. The absence of TRC in the homologous region of *T. varians* genome is consistent with the absence of *Fem* in *T. varians*. It can be inferred that the ancestral transposon of putative origin of satDNA was not active in the lineage of *Trilocha*.

As well as the 5’ half, 3’ half of the HSR was not annotated in the *B*. *mori* genome, but we realized that the W chromosome of *T. varians* harbors significantly similar sequences, and RepeatMasker annotated those sequences as retrotransposon-derived LTR (Fig. 2C; S13 Fig). The inner regions between pairs of this sequence in the W chromosome of *T. varians* encode at least *pol,* occasionally both *gag* and *pol* (S13 Fig), supporting that these sequences are derived from LTR retrotransposon. These interval sequences between LTRs show significant homology to gypsy in *B. mori* (S4 Table). Therefore, we speculated that the 3’ sequence of the *Fem* piRNA-producing region was derived from the LTR of gypsy. If *Fem* piRNA was born out of coincidence, namely the fusion of satDNA and LTR of gypsy, it is consistent with the fact that only two species, namely *B. mori* and *B. mandarina* [19], have been experimentally proven to adopt *Fem* piRNA dependent the sex determination system so far despite approximately one decade having past since the discovery of *Fem*.

## Materials and Methods

### Insects

*B. mori* (p50T strain) was provided from National BioResource Project-Silkworms (NBRP-silkworms; https://shigen.nig.ac.jp/silkwormbase/topAction.do). *T. varians* (NBRP strain, derived from individuals caught in Ishigaki Island, Japan) [20] and *B.* mandarina (Sakado strain) were provided from National BioResource Project-WildSilk (NBRP-WildSilk; http://shigen.nig.ac.jp/wildmoth/). *B*. *mori* and *B. mandarina* larvae were fed on fresh leaves of *Morus alba*, while *T. varians* were fed on fresh leaves of *Ficus microcarpa*. *B*. *mori* and *B. mandarina* were reared under a long-day condition (16 h light / 8 h dark) at 25°C. *T. varians* was reared under a long-day condition (16 h light / 8 h dark) at 20°C.

### Genomic DNA isolation

For the HMW DNA isolation of *B. mori* and *T. varians,* a last-instar female individual immediately after the gut purge and a female pupa 2 days post pupation were used, respectively. The samples were instantly frozen in liquid nitrogen and ground to a fine powder in a mortar. Genomic DNA was isolated using NucleoBond HMW DNA (MACHEREY-NAGEL) according to the manufacturer’s protocol.

### PacBio HiFi library preparation for *B. mori* whole genome sequencing

4 µg of genomic DNA from female *B. mori* was sequenced using the Pacbio SMRT sequencing (HiFi) method. The library was constructed using the SMRTbell Template Prep Kit 3.0 (Pacific Bioscience, USA) and sequencing on PacBio Sequel IIe (NovogeneAIT, Singapore). The library information was summarized in S5 Table.

### *T. varians* whole genome sequencing

For long-read sequencing, 4 µg of genomic DNA from female *T*. *varians* was submitted to the adaptor ligation using Ligation Sequencing Kit SQK-LSK109 (Oxford Nanopore Technologies, UK). The prepared sample was sequenced on Oxford Nanopore GridION (Oxford Nanopore Technologies, UK) using a Flongle cell. For short-read sequencing, the library was prepared using the NEBNext DNA Library Prep Kit (New England BioLabs). The constructed library was sequenced on Novaseq 6000 platform (illumina, USA). The information of each library was summarised in S5 Table.

### Genome assembly of *B. mori* female

HiFiAdapterFilt v 2.0.1 [21] was used for adaptor trimming of Pacbio Hifi reads. The trimmed reads were assembled using hifiasm v.0.16.1-r375 [22]. The output p_ctg.gfa and a_ctg.gfa files were converted to fasta format and merged into a single file. The merged file was aligned against the current male genome assembly (accession number: GCA_014905235.2), then isolated putative W-linked sequences. The W- linked sequences and the remnants (Z-linked and autosomal sequences) were submitted to optical genome mapping for further assembly. The correspondence between scaffolds and chromosomes was summarised in S1 Table.

### Genome assembly of *T. varians* female

Porechop v. 0.2.3 [23] was used for the adaptor trimming of nanopore reads. The Trimmed reads were assembled using flye v 2.9 [24]. The draft assembly was polished by racon v 1.5.0 [25]. Misassembly was corrected using the consensus mode of medaka v 1.7.2 with the “-g” option (https://github.com/nanoporetech/medaka). Error correction using short reads was aligned to the draft assembly for error correction using bwa-mem2 v 2.2.1 [26] and hypo v 1.0.3. (https://github.com/kensung-lab/hypo). This procedure was repeated once again. The error-corrected assembly was submitted to optical genome mapping for further assembly. The correspondence between scaffolds and chromosomes was summarised in S1 Table.

### Optical genome mapping

Genomic DNA was isolated from the pupa immediately after the pupation for the optical genome mapping of the draft genomes of *B. mori* and *T. varians*. DNA isolation was conducted using the Bionano Prep Animal Tissue DNA Isolation Fibrous Tissue Protocol (Bionano Genomics). For *T. varians*, DLE-1 was used for the direct label stain. For *B. mori*, two enzymes, DLE-1 and Nt.BsqQI were used for labelling. The labelling procedure was conducted according to the Bionano Prep Direct Label and Stain Protocol. The labelled samples were scanned on the Bionano Saphyr system using Saphyr Chip G2.3. The obtained data were analysed using Bionano Access v 1.7.1.1 (https://bionano.com/access-software/) and Bionano Solve v. 3.7. (https://bionano.com/software-downloads/).

### Genome assembly assessment by BUSCO and Inspector

To assess the genome assemblies, Inspector [27] and BUSCO v. 5.4.6 with lepidoptera_odb10 lineage dataset were used [28]. The results were summarised in S2 Table.

### Construction of a *T.varians* BAC library

BAC construction was carried out according to a method described in Okumura et al. (2019) with slight modifications [29]. We used male genomic DNAs extracted from *T.varians* pupae (600 mg), and the genomic DNAs were digested with HindlJ (8-12 U/ml) at 37°C for 25 min. The digested fragments were fractionated and collected using CHEF Mapper pulsed field gel electrophoresis (Bio-Rad). The extracted DNA fragments were ligated to the pBeloBAC11 vector, and the ligates were transformed by electroporation (GenePulser II, Bio-Rad) into DH5α Electro-Cells (TaKaRa). The electroporated cells were spread on LB plates containing 12.5 mg/l chloramphenicol (Cm), X-gal and isopropyl β-D-thiogalactopyranoside. Grown white colonies were stocked with 384-well plates. Stocked plates were stored at ™80°C until further use.

### Chromosome preparations

Chromosome specimens were prepared according to a method described in Yoshido et al. (2014) [30]. Briefly, ovaries and testes of the last instar larvae of *B. mori* and *T. varians* were dissected in a physiological solution, and testes and ovaries were treated with 75 mM and 100 mM KCl solution for 15 min, respectively. After the hypotonic treatment, the testes and ovaries were fixed in Carnoy’s (ethanol: chloroform: acetic acid, 6:3:1) for 10 min. Spermatocytes and oocytes were transferred into a drop of 60% acetic acid on a slide glass and spread at 55°C using a heating plate. The preparations were passed through 70%, 80% and 99% ethanol series, air dried and stored at ™20°C until time to use.

### BAC-FISH mapping

According to the newly constructed genome assembly data of *T. varians* and *B. mori*, we designed *T. varians* STS primers for the genomic positions corresponding to two opposite positions in each *B. mori* chromosome (S7 Table). We also designed 3 *B. mori* STS primer pairs from the female-specific sequences to select the W BACs (S7 Table). Using the STS primer pairs, we selected the BACs according to the methods written in Yoshido et al. (2014) [30]. The PCR-selected BACs were cultured in 1.5 ml of LB solution containing 20 mg/l Cm for 16 h at 37 LJ with a shaking incubator (Bio Shaker BR-23FH, Taitec). Then, plasmid DNA was extracted with a standard alkaline SDS method. BAC-DNA Probe labelling and BAC-FISH were performed according to a method described in Yoshido et al. (2014) [30]. The FISH preparations were counterstained and mounted with Vectashield Antifade Mounting Medium with DAPI (Vector Laboratories). A Leica DM6000B fluorescence microscope (Leica Microsystems) and a DFC350FX black and white charge-coupled device camera (Leica Microsystems) were used for observation and image capturing. The images were processed with Adobe Photoshop 2022.

### W-linked *Fem* mapping

FISH mapping of *B. mori* Fem clusters was done by simultaneous Z-chromosome identification with 3 Z- BAC-FISH. Briefly, the Z chromosome thread of WZ bivalents in the female pachytene nucleus is identified with 9A5H, 14I7D and 5H3E BCA-probes [31]. Because the Z pairing partner is the W chromosome, hybridisation signals from cloned *Fem* probe [32] appear on the W thread. The positions of *Fem* probe signals on the W chromosome can be identified relative to the three BAC positions mapped on the Z chromosome. Repetitive elements annotation in the genomes Repetitive elements in the genome assemblies were identified using RepeatModeler v 2.0.4. [33]. Then, the annotated elements were masked using RepeatMasker v 4.1.2. (http://www.repeatmasker.org).

### Chromosome-scale genome comparison

For nucleotide sequence comparison, Mummer v. 4.0.0 [34] was used. For the synteny analysis between genome assemblies of *B. mori* and *T. varians*, genomic coordinate of single copy orthologs predicted by BUSCO analysis were used. The synteny plot was created by MCScanX [35]

### RNA sample preparation for piRNA-seq, RNA-seq, Iso-seq and ATAC-seq

Total RNA was extracted from multiple early embryos using TRIzol reagent (Invitrogen) according to the manufacturer’s protocol. For *B. mori* and *B. mandarina*, embryos were sampled at 24 hours post oviposition, while *T*. *varians* embryos were sampled at 72 hours post oviposition. The piRNA-seq library was prepared using TruSeq small RNA kit (illumina) according to the manufacturer’s protocol with a slight modification. To sequence piRNA, a region of 147-158 nucleotides was extracted in the purification step of the cDNA construct using BluePippin (Sage Science, USA). The constructed library was sequenced on illumina Hiseq 2500 platform (illumina, USA). RNA-seq library was prepared using NEBNext Poly(A) mRNA Magnetic Isolation Module (New England BioLabs) and NEBNext® Ultra™ ll Directional RNA Library Prep Kit (New England BioLabs) according to the manufacturer’s protocol. The constructed library was sequenced on the illumina Novaseq6000 platform (illumina, USA). For Iso-seq, the library was constructed using Sequel Iso-seq Express Template Prep (Pacific Bioscience, USA) according to the manufacturer’s protocol. The constructed library was sequenced on PacBio Sequel platform (PacBio, USA). Fragmentation and amplification of the ATAC-seq libraries were conducted according to Buenrostro et al. (2015) [36]. The constructed libraries were sequenced on the Illumina HiSeq.

### Transcriptome data analysis

Fastp v 0.20.1 [37] was used for the quality trimming of short-read data. Iso-seq subreads were converted to circular consensus sequences (ccs) using ccs v 6.4.0. The resulting ccs reads were assembled with the trimmed short reads using rnaSPAdes v 3.15.5 [38]. The resulting transcripts.fasta file was aligned to the corresponding genome data using pasa v 2.5.2 [39]. For *B. mandarina*, pasa alignment was not performed due to the absence of female genome data. The libraries’ information was summarized in S7 Table.

### Collinearity detection between the W chromosome and Z chromosome

From the transcriptome data of early embryos of *T. varians*, the genes were extracted which met the following criteria:

1. not a transposon or a transposable element-related.
2. transcribed from the Z or W chromosome.
3. the top two alignments yielded by BLASTN search are the chromosome to which the gene is linked and its homologous chromosome, respectively, i.e., we searched for genes (transcripts) which are Z- linked ones whose putative homologues were W-linked and which are W-linked ones whose putative homologues were Z-linked.

### Small RNA mapping

The small RNA reads were trimmed using Trim Galore v 0.6.6 (https://github.com/FelixKrueger/TrimGalore) in small RNA mode. The trimmed small RNA reads were mapped to the assembled transcriptome, allowing up to 3 nucleotide mismatches using Hisat2 v 2.1.0. [40] and ngsutils v 0.5.9. [41]. The infomation of each library was summarized in S8 Table.

### In silico piRNA cluster analysis

To identify the piRNA cluster, proTRAC v. 2.4.4 [42] was used with following options: ‘-clsize 5000 - pimin 23 −pimax 29 ™1Tor10A 0.3 ™1Tand10A 0.3-clstrand 0.0-clsplit 1.0-distr 1.0-99.0-spike 90-1000-nomotif-pdens 0.05’

### Visualisation of tandem repeat structures

To identify and visualise tandem repeat structures on chromosome 8 of *B. mori*, *B. mandarina* and *T. varians*, StainedGlass v. 0.5 was used [43]. The bin size of the window in which to breakup the input sequence all by all alignment was set to “200.”

### Locating *Fem*, satellite DNA-like sequence, and gypsy-like sequence

To identify *Fem* copies with full length, BLASTN search v 2.9.0 [44] was conducted with “-perc_identity 90-qcov_hsp_perc 90” options. NCBI genbank accession number AB840787 was used as *Fem* sequence. In the BLASTN search of the satDNA-like and Gipsy-like sequences, BLASTN search was performed with the “-task blastn_short” option since the lengths of satDNA-like and gypsy-like were less than 50 nucleotides. The following sequences were used as queries of satDNA-like and gypsy-like sequences, respectively: “TTTCTTTCATTGTTACCTCTTTTTG” and “TGTCAATTCATAAAGTCATTCAGTGT.” To identify BovB-like sequences, a BLASTN search was conducted with the default option. NCBI Genbank accession number LC763122.1 was used as a query. Multiple alignments of gypsy, satDNA-like sequence and BovB were performed by mafft v 7.520 [45].

## Acknowledgments

Insects were donated from Kyushu University and Shinshu University according to a Grant-in Aid “National Bio Resource Project (NBRP, RR2002), Silkworm Genetic Resources” for Scientific Research from the Ministry of Education, Science, Sports and Culture of Japan. The optical genome mapping was performed by AS ONE CORPORATION. We would like to thank Dr. Hirohisa Kishino for his precise comments on the manuscripts.

## Author Contributions

J.L. designed the research plan, performed genomic DNA and RNA extraction, analyzed the obtained data, and wrote the manuscript. T.S. also designed this research plan and performed the part of data analysis. T.F. and K.S. performed BAC-clone selection and chromosome imaging. A.T. also designed the research plan and performed preliminary experiments. K.Y. and S.S. performed genomic DNA extraction nanopore GridION operation.

## Competing Interest Statement

The authors declare that they have no competing interests.

## Data availability

Supplementary information is available in the online version of the paper. The deep-sequencing data from this study have been deposited in the DDBJ Sequence Read Archive (SRA) database. Genomic data of *B. mori* and *T. varians* females were under the accession code PRJDB15448. piRNA-seq, RNA-seq, and Iso-seq data were under the accession code PRJDB13955. The accession numbers of sequences comprising female genome assemblies of *B. mori* and *T. varians*were AP028171–AP028199 and AP028256– AP028282, respectively. We used the genome assembly of *B. mandarina,* which was deposited in NCBI under the accession numbers AP028283– AP028309. The assembled transcriptome data and output files of RepeatModeler and RepeatMasker used for analysis can be downloaded at 10.6084/m9.figshare.22657477.

## Notes

### Competing Interest Statement

The authors have declared no competing interest.

