## Supplementary material for "Transposon fusion gave birth to *Fem*, *Bombyx mori* female determining gene": S11 Fig

**a**

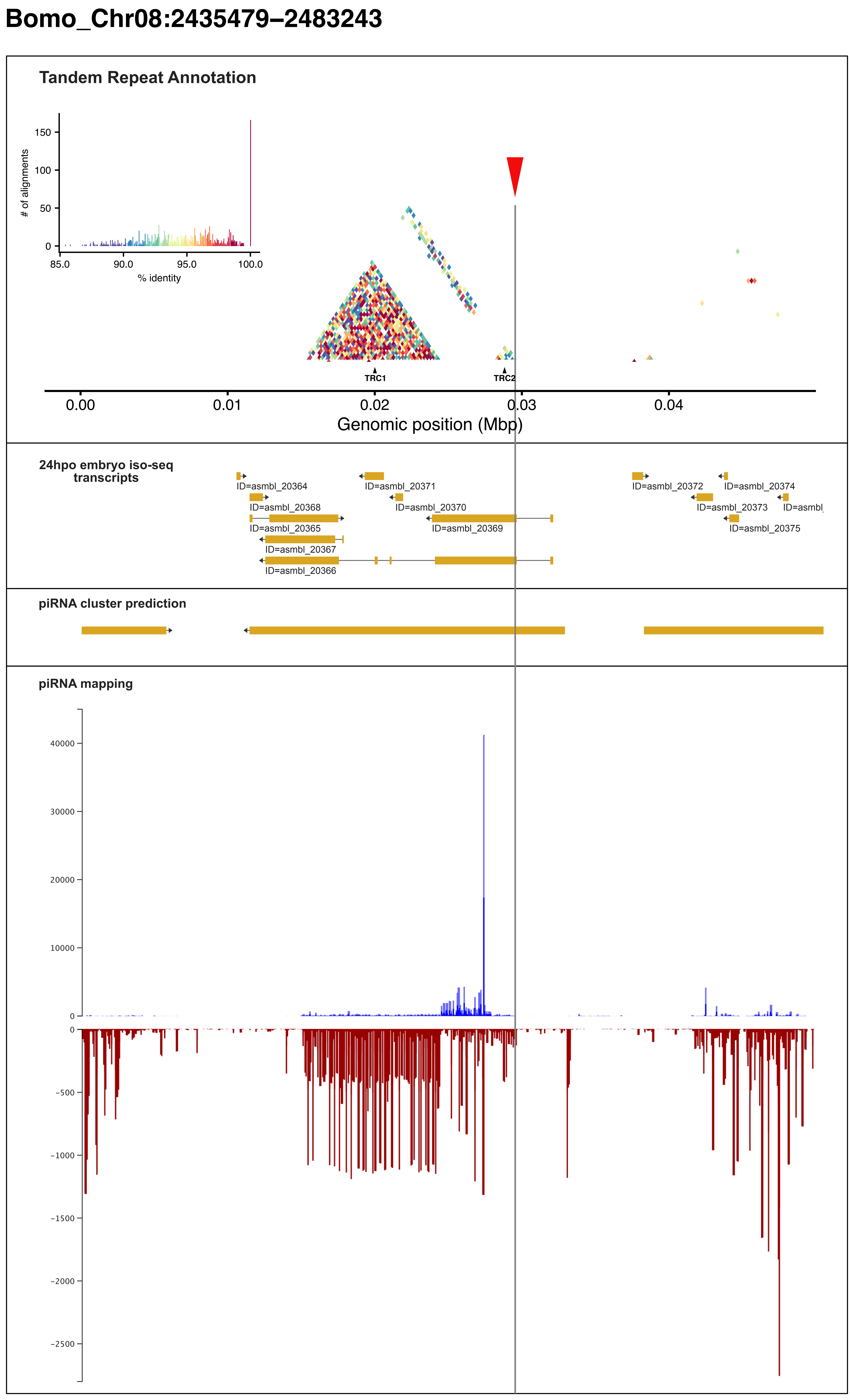


**b**

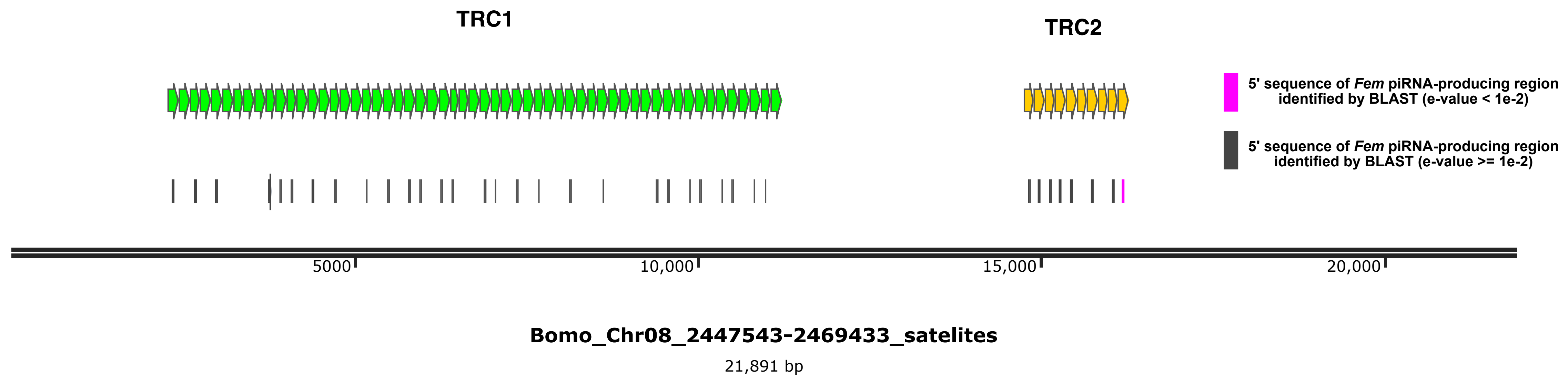


**c**

```
motif1        nnnnnnacttctttttcaaacattgcttattaagactggcnattttagcagtcaagtgtg
motif2        ------acttcctcttcaaacattgcttattaagactggcaattttagcagtcaagtgtg
                    *****.*.************************** ******************

motif1        attncnttctattgtnntacctctttgtggatcttcgctgcagactgttgtgaatctacn
motif2        anttgcttccattgt--tacctctttgtggatcttcgctgcagactgttgtgaatctacc
              * *   ***.*****  ****************************************

motif1        tccgtaagatatttatattgttat---------gttntatagtttctaaggtattcagnn
motif2        tccgtaagnnnnnnnnnnnnatattcatatgnnnttatatagtttctaaggtattcag--
              ********              ***         ** ********************

motif1        nnnnnnnnnnnnnnnnnnnnnnnnn
motif2        -------------------------
```

**d**

```
1       acttcctcttcaaacattgcttattaaaactggcaattttagcagtcaagtgtga-ttgc
2       acttccacttcaaacattgcttattaagactggcaattttagcagtcaagtctga-ttgc
3       acttcctcttccaacattgcttattaagactggcaattttagcagtcacgtgtga-ttgc
4       acttcctcttccaacattgcttattaagactggcaattttagcagtcacgtgtga-ttgc
5       atatccacttcaaacattgcttattaagactggcaattttagcagtcaagtctca-ttgc
6       acttccacttcaaacattgcttattaagactggcaatttgagcagtcaagtgtgacttct
7       acttcctcttcaaatattgcttgttaaaactggcgaccttaacagtcaagtgtgt-ttac
8       aattcctcttcaaacattgcttattaagaatggcagttttagcagtc-attgtga-ctcc
9       acttcctcttcaaacatagcttattaagactggcaattttagcagtcaagtgtga-ttgc
10      acttcctcttcaaacatagcttattaagactggcaattttagcagtcaagtgtga-cttc
        *  *** **** **.** ****.****.* ****....* *.*****   * *   .*  .
```

5' sequence of *Fem* piRNA-producing region ► tttc

```
1       ttccattgttacctctttatggatcttcgctgcggactgttgtgattctacctccgtaag
2       ttccattgttacctctttatggatcttcgttgcagactgttctgaatctatctccgtaag
3       ttccattgttacctctttatggatcttcgttgcagactgttctgaatctatctccgtaag
4       ttccattgttacctctttgtggatcttcgctgcggactgttgtgattctacctccgtaag
5       ttccattgttacctctttgtggattttcgctgcagactgttgtgaatctacctccgtaag
6       ttttattgttacctctttgtggatcttcgctgcagactgttgtgaatctacctccgtaag
7       ttccattgttacctctttgtggatcttcgctgcagtctgctgtgaatctacttccgtaag
8       ttctattgttacctctttgtggatctttgctatacactattgtgaatct-----------
9       ttccattgttacctctttgtggatctttgctgcacactgttgtgaatct-----------
10      ttccattgttacctcttttgggatcttcgctggagactgttgtgaagctattcccgttac
        **..************** *****.**.*.*. .  **..* ***   **
```

► tttcattgttacctcttttg

```
1       ------------atattcatatg---ttatatagtttctaaggtattcaa
2       ------------atattcatatg---ttatatactttctaaggtactcag
3       ------------atattcatatg---ttatatactttctaaggtactcag
4       ------------atattcatatg---ttgtatagtttctaaagaattcag
5       ------------atattcatatg---ttatatactttctaaggtattcag
6       ------------atattcatatg---ttgtataatgtctaaagtattcag
7       ctataagtatttatatcgttatc---ttatatagtttctaaggtattcag
8       ------------atatttatatg---ttgtacagtttttaaggtattcag
9       ------------atatttatatg---ttgtatagtttctaaggtattcag
10      ------------tttatgatgcaagcccggccgctgtcgaacctgttcag
                     *  .  *..    ...  ..  * *. **   ..***.
```
